# Experimental Evolution of Insect Immune Memory VS. Pathogen Resistance

**DOI:** 10.1101/137703

**Authors:** Imroze Khan, Arun Prakash, Deepa Agashe

## Abstract

Under strong pathogen pressure, insects often evolve resistance to infection. Many insects are also protected via immune memory (‘immune priming’), whereby sub-lethal exposure to a pathogen enhances survival after secondary infection. To understand the evolution and consequences of these immune responses, we imposed strong pathogen selection on flour beetles (*Tribolium castaneum*), infecting them with *Bacillus thuringiensis* (Bt) for 11 generations. Populations injected first with heat-killed and then live Bt each generation evolved high basal resistance against multiple Bt strains. In contrast, all populations injected only with a high dose of live Bt evolved less effective but strain-specific priming response. Control populations injected with heat-killed Bt did not evolve priming; and in the ancestor, priming was effective only against a low Bt dose. Thus, pathogens can select for rapid modulation of insect priming ability, leading to divergent immune strategies (generalized resistance vs. specific immune priming) with distinct mechanisms and adaptive benefits.

## INTRODUCTION

A large body of work shows that strong pathogen pressure drives the evolution of resistance mechanisms in insect hosts, reducing the fitness impact of infection (Fellowes *et al.* 1999; Kraaijeveld & Godfray 2008; Boots & Begon 2009; Vijendravarma *et al.* 2009; Faria *et al.* 2015; Gupta *et al.* 2016). In addition, many insects exhibit a form of immune memory (priming response), gaining increased protection against a pathogen after initial exposure to a low dose of infection (reviewed in Contreras-Garduño *et al.* 2016; Milutinović *et al.* 2016). Immune priming is also observed across multiple natural populations of flour beetles (Khan *et al.* 2016). Together, this work suggests that in addition to basal resistance, immune priming is a significant immune strategy across insects. Mathematical models predict that such immune memory can reduce the impact of disease prevalence (Tate & Rudolf 2012) and alter population dynamics (Tidbury *et al.* 2012). However, we have limited empirical information about the evolutionary benefits of priming relative to the innate immune responses that confer basal resistance against a pathogen (without priming) in insects.

We also know very little about the selective pressures and ecological conditions that shape the evolution of the insect immune priming response vs. basal resistance to a pathogen. A general mathematical model examining the impact of various immune strategies on population growth suggests that the frequency and duration of infection are key determinants of immune function (Mayer *et al.* 2016). Although this model did not specifically address the evolution of immune memory vs. basal resistance in insects, it makes the general prediction that adaptive immunity (responsible for immune memory) is more likely to evolve when infection is relatively rare. In contrast, assuming greater maintenance costs of constitutively expressed resistance, innate immunity is predicted to be more advantageous under frequent infection. However, it is unclear whether this assumption holds for insects, because we have limited information about the molecular mechanisms responsible for insect immune priming. Recent evidence suggests that components of innate immunity – cellular and humoral defenses – play a critical role in the priming response of Dipterans (Pham *et al.* 2007;

Rodrigues *et al.* 2012; Weavers *et al.* 2016). However, the extent of overlap between priming and resistance pathways, and their relative costs, are not well understood. Thus, from an evolutionary perspective it is not even clear whether insect immune memory and resistance are distinct phenomena. In contrast to direct resistance, we do not know whether priming can evolve rapidly, and whether (and when) it is adaptive in natural populations.

To begin to understand the impact of pathogen pressure on the evolution of alternative immune strategies (priming vs. resistance), we allowed replicate outbred laboratory populations of the flour beetle *Tribolium castaneum* to evolve with their natural pathogen *Bacillus thuringiensis* (strain DSM 2046, henceforth Bt, isolated from a Mediterranean flour moth as described in Roth *et al.* 2009; Khan *et al.* 2016). The pathogen imposes significant mortality on the ancestral beetle population (~65% mortality within 2 days after infection; Figure S1), imposing strong selection on their immune function. Importantly, although the ancestral population was capable of mounting a priming response against Bt infection (~8000 cells per beetle; Khan *et al.* 2016), this basal priming ability was ineffective against the relatively higher dose of infection used here (~12000 cells; Figure S1). Thus, at the beginning of the experiment, beetle populations had low effective basal resistance as well as no priming ability against Bt. We tested whether populations evolve stronger priming or higher resistance when exposed to a single severe infection each generation (no priming opportunity) vs. when given the opportunity for priming (first injected with heat-killed bacteria, then infected with live pathogens). We found that populations showed divergent responses to Bt infection, either evolving immune priming against the specific Bt strain used for selection, or more effective but non-specific basal resistance.

## RESULTS

We allowed replicate beetle populations to evolve under different selection regimes designed to impose strong pathogen selection (infection with live Bt cells each generation; Figure 1A, see Methods for details). Populations evolved with or without an opportunity for priming (pricked with buffer or heat-killed Bt cells before live Bt infection). Control populations evolved without direct selection by Bt (Figure 1A). After 8 and 11 generations of selection, we propagated parallel populations under relaxed selection for two generations, and thereafter measured priming ability (survival rate of unprimed vs. primed beetles) and basal resistance (survival rate of unprimed vs. uninfected control beetles) to Bt in these standardized populations (see Figure 1B). As expected, the high infection dose in each generation imposed substantial mortality (and therefore strong selection) on populations in I (infection only) and PI (priming + infection) regimes. However, within a single generation of selection, post-infection survival (after 2 days) increased from ~40% in the ancestor to ~45% in I populations, and as high as ~68% in PI populations (Figure 2). After 11 generations of selection, adult survival increased further: ~50% adults in I populations and ~80% adults from PI populations survived infection. As expected, adults in the C (control) and P (priming only) populations that were never exposed to Bt maintained very high survival (~90-100%) throughout the experiment (Figure 2). Note that regardless of variation in post-infection survival, we used 60 mating pairs to initiate each successive generation for all populations (accounting for expected mortality, we infected a larger number of beetles; see Methods). This design allowed us to ensure strong selection on survival every generation without imposing a bottleneck.

**Figure 1.**
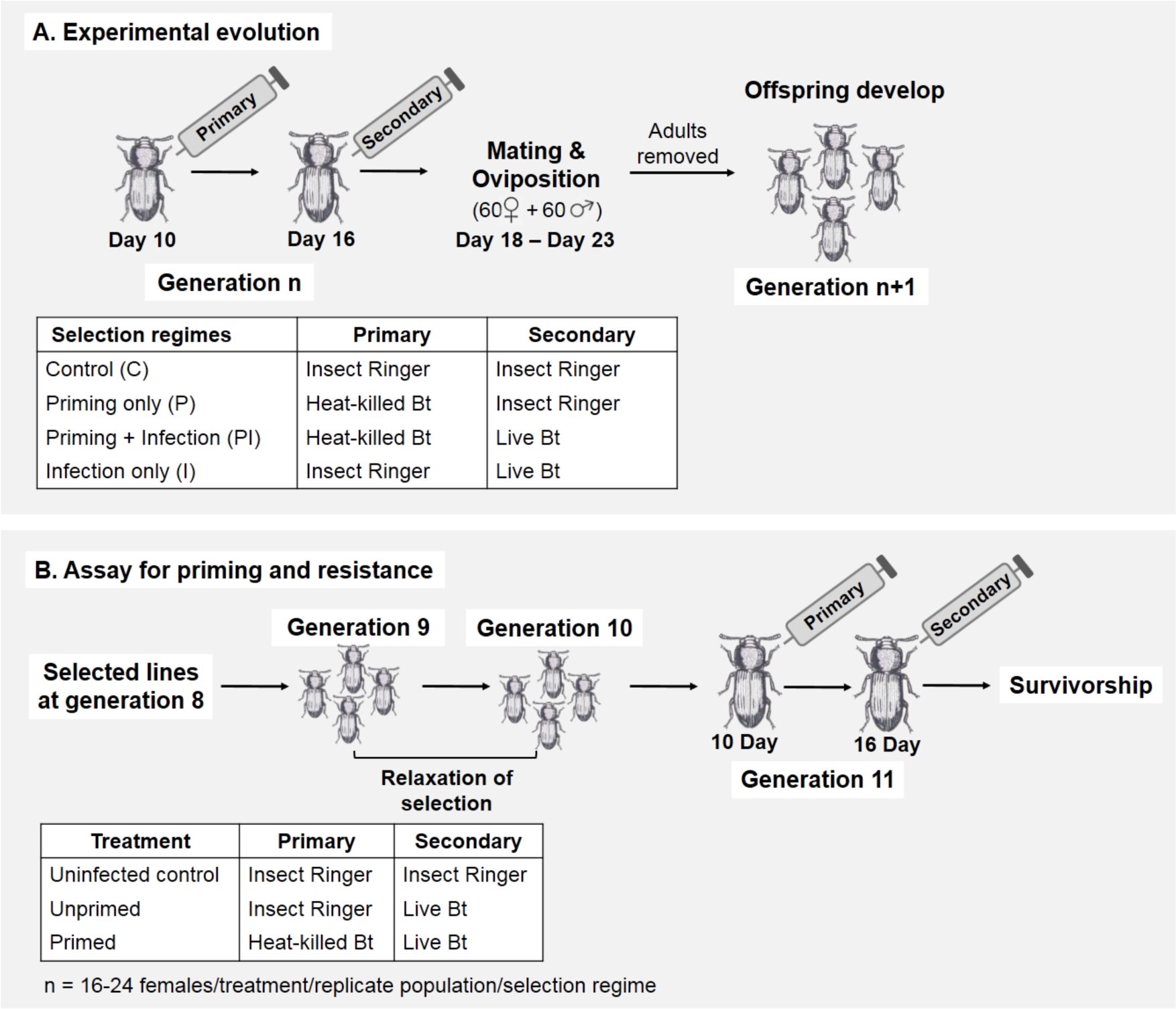
(A) Design of experimental evolution and selection regimes. Each selection regime had 4 replicate populations. All PI populations were first injected with heat-killed and then live *B. thuringiensis* (DSM 2046; Bt) each generation, whereas all I populations were directly exposed to live Bt each generation without an opportunity for priming. C and P populations (controls) were never exposed to live Bt. (B) Generating standardized beetles to measure evolved priming and resistance. After 8 generations of pathogen selection, we set up parallel populations that experienced relaxed pathogen selection for two generations. For each such “standardized” population, we compared survival of unprimed vs. primed and unprimed vs. uninfected control beetles to estimate priming response and resistance to infection respectively. We followed a similar protocol to generate standardized beetles after 11 generations of pathogen selection.

**Figure 2.**
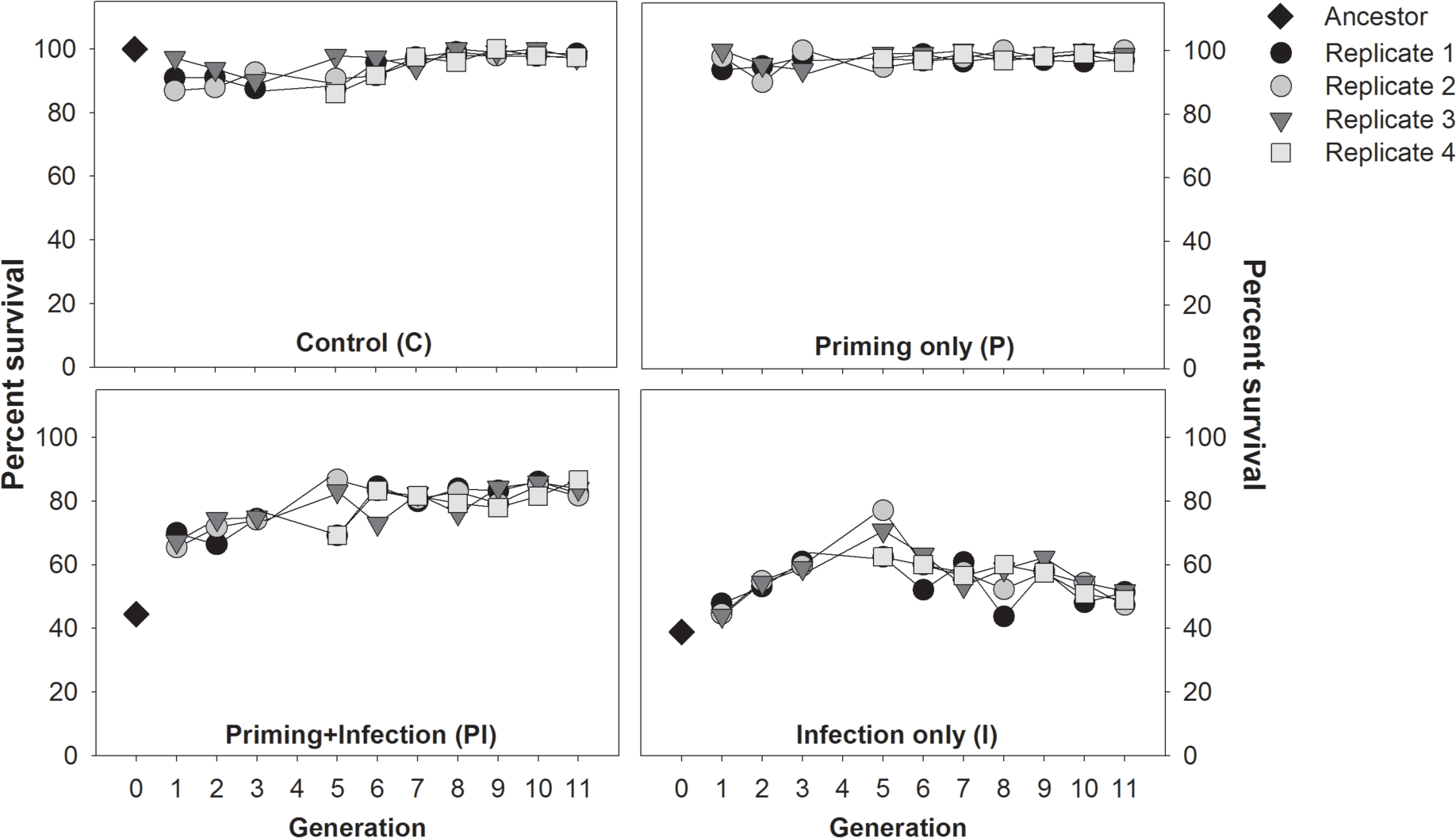
Adult survival during the first 48 hours after secondary infection with live *B. thuringiensis* (DSM 2046) cells (day 16-18; see Figure 1A), during the course of experimental evolution. 60 pairs of survivors from each population were allowed to mate and produce the next generation.

To determine the mechanism responsible for decreased post-infection mortality in evolved PI and I regimes, we used females from standardized populations generated after 8 generations of selection. As expected, females from populations that were not exposed to live pathogen (unhandled ancestral population, and populations from C and P regimes) showed high mortality after infection and no survival impact of priming (Figures 3A-C and first two panels of Figures 3F and 3G). Thus, neither priming nor resistance had evolved in these populations. We found that highly effective basal resistance to Bt had evolved in 3 of 4 PI populations, conferring survival rates nearly as high as uninfected control beetles (Figures 3D, 3^rd^ panel of Figure 3G). However, in the fourth population (population PI4), we observed significant priming ability that conferred a 2-fold survival advantage (3^rd^ panel of Figure 3F). Note that all PI populations were first injected with heat-killed and then live Bt each generation, allowing an opportunity for the evolution of priming-induced survival benefits. Despite this opportunity for priming, only one population showed evidence of priming ability after 8 generations of selection. In contrast to PI populations, 3 of 4 populations in the I regime (infected with a single high dose of Bt) evolved priming ability rather than resistance. Although unprimed I beetles remained highly susceptible to infection, priming resulted in significantly improved survival (~3-fold increase in survival, Figure 3E, 4^th^ panel of Figures 3F and G). Interestingly, the fourth population (I3) – despite similar survival as other replicate populations during experimental evolution (compare I3 with I1, I2 & I4; Figure 2) – evolved neither priming nor improved basal resistance (4^th^ panels of Figure 3F & G).

**Figure 3.**
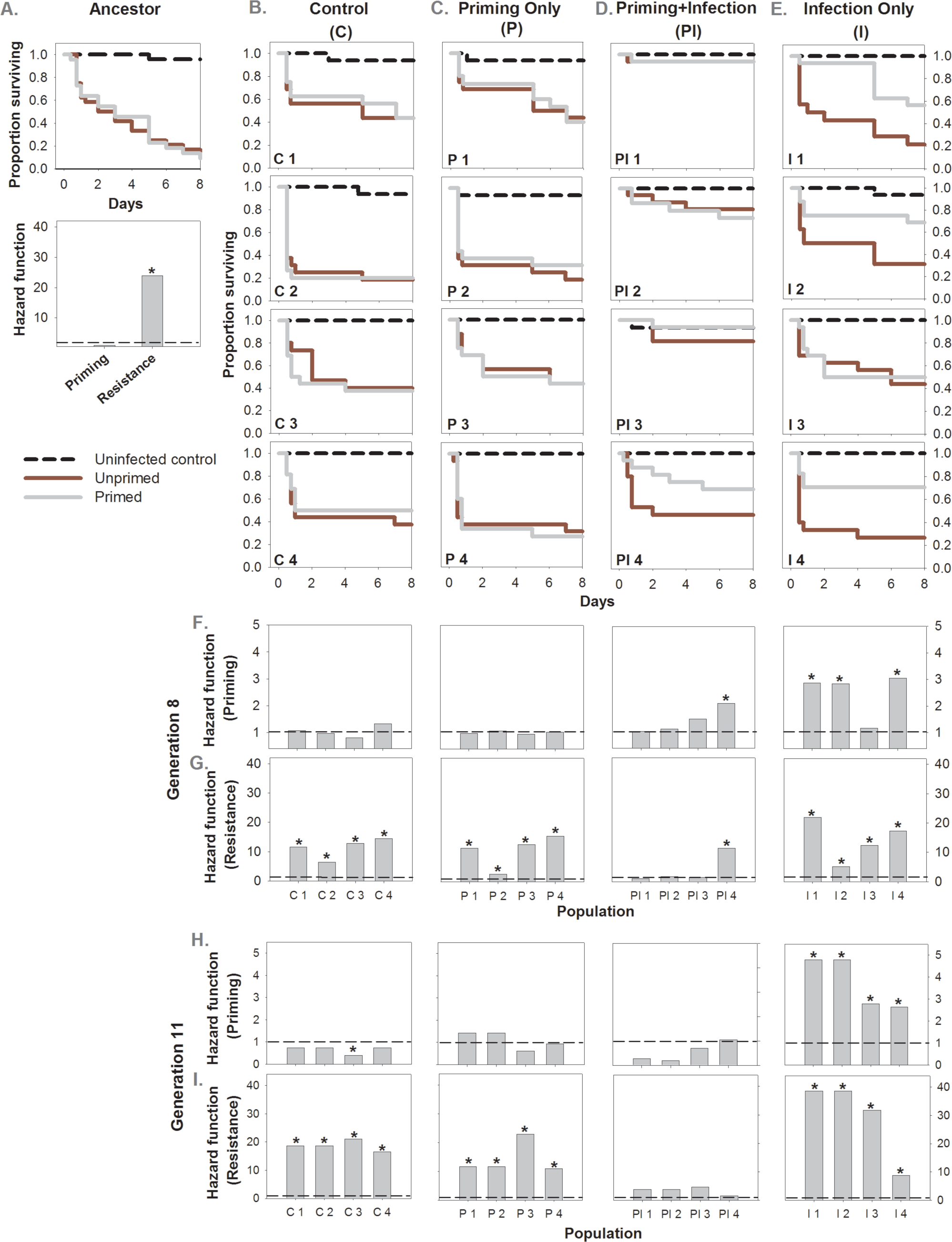
(A) Survival curves and associated hazard ratios for ancestral population as a function of bacterial infection and priming (n=22-24 females/treatment). Evolution of priming vs. basal resistance in response to selection imposed by *B. thuringiensis* DSM 2046 (Bt), after 8 generations (B-G) (*n* = 16-24 females/treatment/replicate population/selection regime) and after 11 generations of selection (H-I) (*n* = 16-26 sex/treatment/replicate population/selection regime). The impact of bacterial infection was calculated as the estimated hazard ratio of deaths occurring in the unprimed infected group compared to the uninfected control group. A greater hazard ratio indicates higher susceptibility to infection, or lower pathogen resistance. Survival benefit of priming response was calculated as the hazard ratio of the proportion of deaths occurring in unprimed vs. primed beetles under a proportional hazard model; greater hazard ratio indicates higher benefit of priming. Survival curves for 4 replicate populations of (B) Control (C) Priming only (D) Priming and Infection (E) Infection only regimes as a function of Bt infection and priming. Estimated hazard ratio for (F, H) Survival benefit of priming and (G, I) Resistance to infection. Horizontal dashed lines in panels F-I indicate a hazard ratio of 1. Asterisks in panels F and H denote significant (p ≤.05) impact of immune priming (i.e. primed beetles survived better than unprimed beetles). Asterisks in panels G and I denote a significant (p ≤.05) susceptibility to bacterial infection compared to uninfected controls (i.e. a lack of resistance against Bt).

We found comparable results when we analyzed female survival beyond the experimental selection window (until day 50): all replicate populations from the I regime showed a 2- to 4- fold survival benefit of priming, and 3 of 4 PI populations showed increased basal resistance with similar lifespan to control females but no priming ability (see Figure S2C and D for survival curves; see Figures S2E and F for hazard ratios). The only replicate population (PI4) that did not evolve higher resistance showed priming instead (2-fold survival benefit; Figure S2C, E-F). Analyzing standardized females derived after 11 generations of pathogen selection, we found that females from all I populations now showed priming (2-4-fold survival benefit) within the selection window (Figure 3H), and all PI populations showed higher resistance (Figure 4I; see Figure S3 for survival curves). Thus, females from population PI4 showed priming after 8 generations of selection, but subsequently evolved resistance within 3 additional generations (compare 4^th^ panels of Figures 3F & 3I). For three replicate populations from each selection regime, we also measured priming and resistance in standardized males after 11 generations of selection. Similar to females, males from three PI populations evolved resistance, and 2 of 3 I populations evolved significant priming response (~4-fold survival benefit) (see Figures S4C and D for survival curves; see Figures S4E and F for hazard ratios). The third I population showed a similar trend (~2-fold survival benefit of priming), but the response was only marginally significant (p = 0.061; see Figures S4D and E). Overall, it thus appears that both sexes evolve similar immune strategies in response to pathogen selection (compare Figure S3 and S4). Together, our results suggest that under pathogen selection, a population can either evolve improved resistance or priming ability but not both. Importantly, our results also suggest that the survival benefit of evolved basal resistance or priming lasts beyond the selection window.

**Figure 4.**
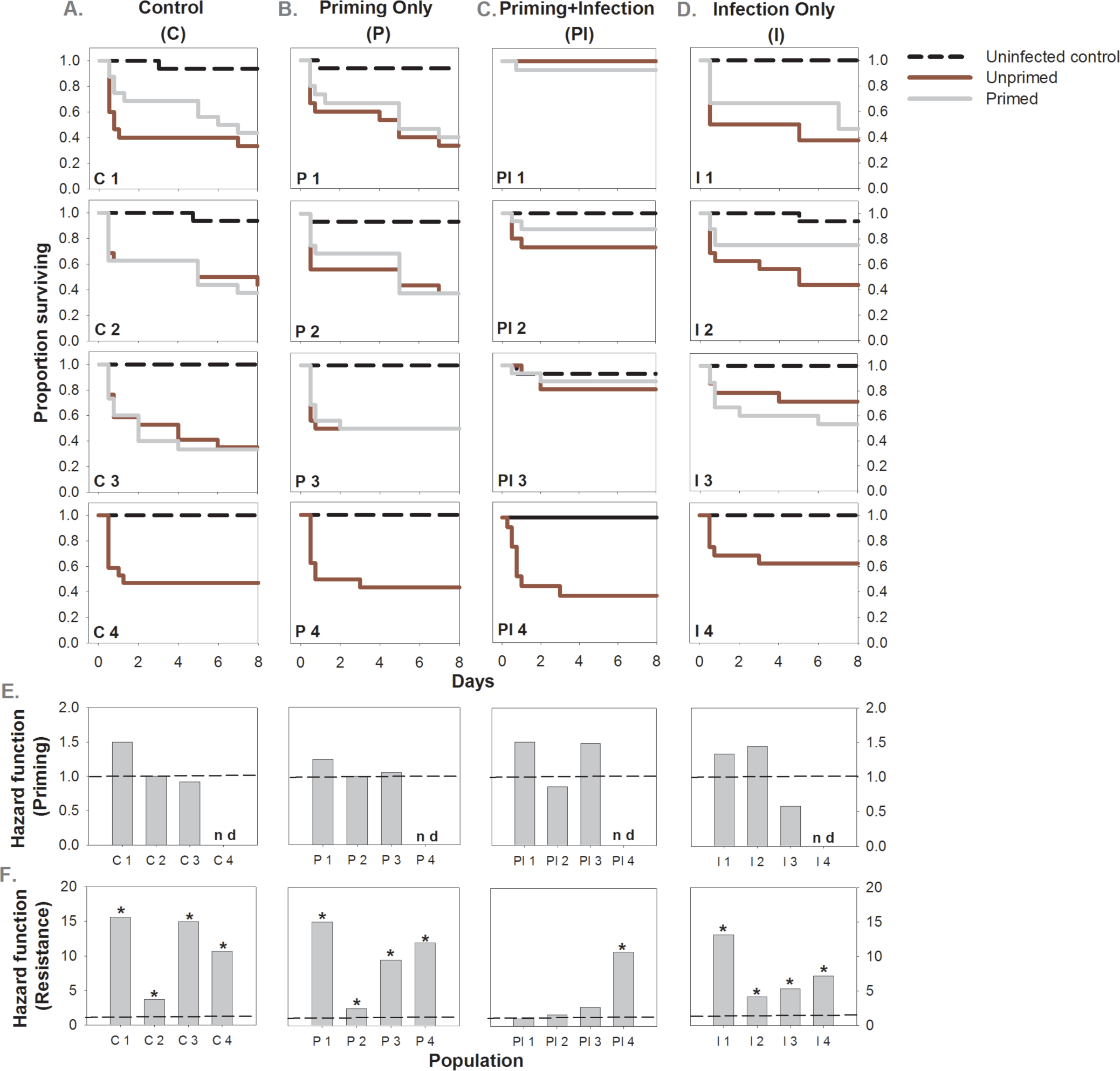
Survival curves for replicate populations of (A) Control (B) Priming only (C) Priming and Infection and (D) Infection only regimes, as a function of *B. thuringiensis* 6905 (Bt1) infection and priming (*n* = 16-24 females/treatment/replicate population/selection regime). Estimated hazard ratio for (E) survival benefit of priming and (F) resistance to infection. We calculated hazard ratios as described in Figure 3. The hazard ratio for priming was not determined for populations C4, P4, PI4 and I4 (denoted ‘nd’ in panel E). Horizontal dashed lines in panel E and F indicate a hazard ratio of 1. Asterisks in panel F denote a significant (p ≤ 0.05) survival impact of bacterial infection, indicating the lack of resistance to Bt.

We used standardized females derived at generation 8 to also test whether the evolved priming response and resistance were specific to the Bt strain used to impose selection. We found that none of the populations showed priming ability against another highly virulent *B. thuringiensis* strain (MTCC 6905, henceforth Bt1; see Figures 4A-D for survival curves and Figure E for hazard ratios; see Methods for details). Thus, the evolved priming response was specific to the pathogen strain imposing selection. In contrast, we found that evolved resistance was non-specific: PI populations that evolved resistance against Bt were also more resistant to Bt1 (3 of 4 populations, except PI4; see Figure 4C for survival curves & Figure 4F for hazard ratios). Finally, we tested whether the evolved survival benefit against Bt infection requires specific priming with the same strain (homologous priming). We found that only beetles receiving homologous priming showed a survival benefit (2.5 to 3-fold survival benefit; Figures 5B & C), whereas heterologous priming (priming with Bt1) failed to protect against Bt infection (Figures 5B & D). Thus, the evolved priming in I populations requires strain-specific immune activation by Bt, rather than a general non-specific immune induction.

**Figure 5.**
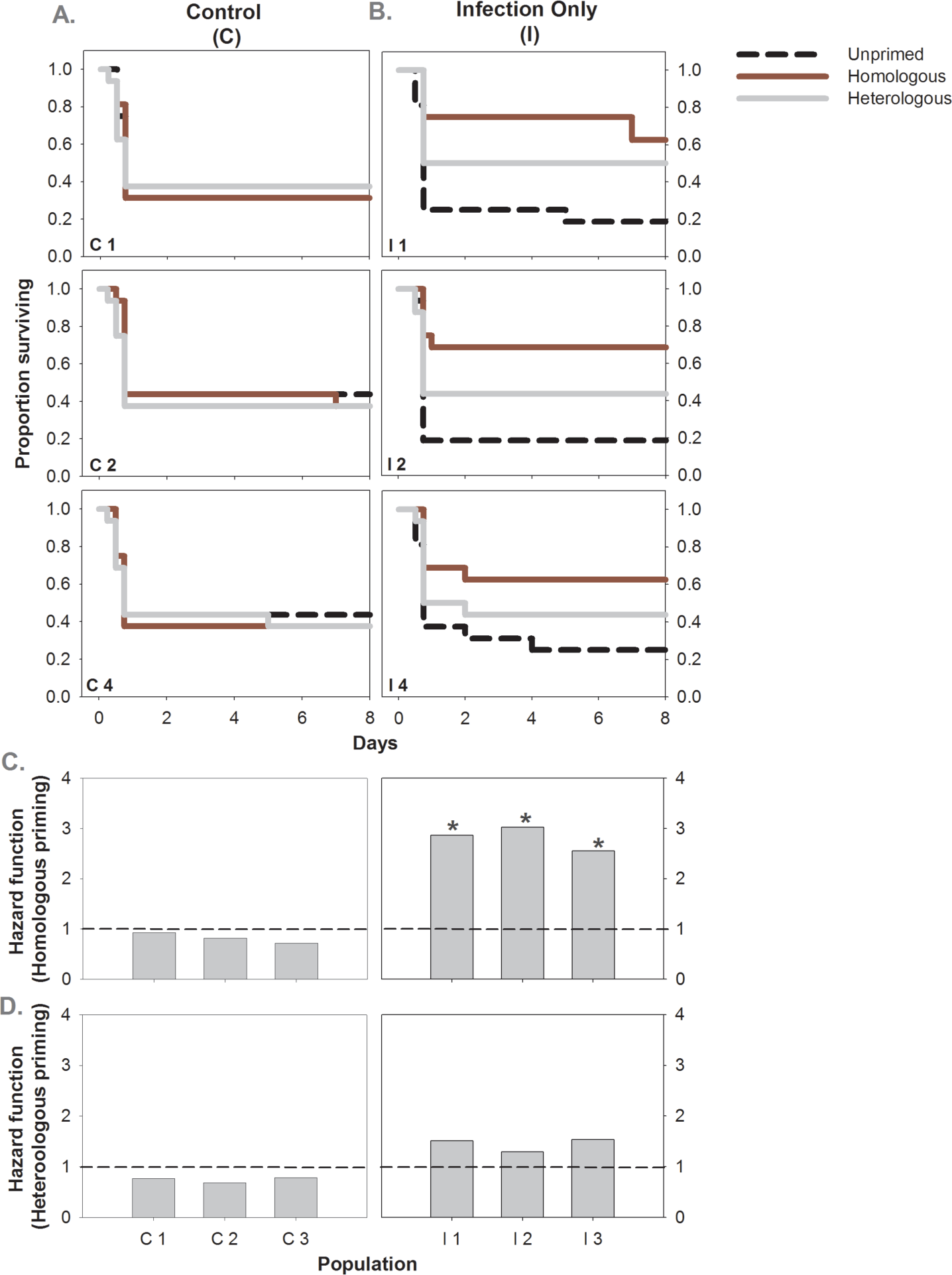
Survival benefit as a function of homologous priming (priming and challenge with *B. thuringiensis* DSM 2046, Bt) vs. heterologous priming (priming with *B. thuringiensis* MTCC 6905, Bt1 and challenge with Bt) in (A) control and (B) infection only lines (*n* = 16-24 females/treatment/replicate population/selection regime). Estimated hazard function for the (C) impact of homologous and (D) heterologous priming in control and infection only lines. We calculated hazard ratios as described in Figure 3. Horizontal dashed lines in panel C and D indicate a hazard ratio of 1. Asterisks in panel C denote a significant (p ≤ 0.05) impact of immune priming, indicating greater survival of primed beetles.

## DISCUSSION

Here, we used experimental evolution to establish the causal link between pathogen-imposed selection and rapid, adaptive evolution of immune memory in an insect. We found that priming ability evolved repeatedly in I populations, where beetles were directly exposed to a single high dose of infection with *B. thuringiensis* (strain DSM 2046; Bt) each generation. Importantly, beetles from these populations were as susceptible to the pathogen as control populations (C & P regimes), and showed no increase in basal resistance. Thus, the evolved priming ability did not confer substantial survival benefits during experimentally imposed selection. Surprisingly, we found that despite an opportunity for priming, PI populations consistently evolved increased basal resistance rather than significant immune priming. Thus, immune priming and improved basal resistance seem to be mutually exclusive responses to pathogen pressure. Intriguingly, one population (PI4) first evolved priming ability before gaining resistance, suggesting that immune priming may be selectively favored as an intermediate step preceding increased resistance. Although this single observation should be interpreted cautiously, it suggests the potential for rapid and complex dynamics in the evolution of alternate immune strategies.

We found that the evolved priming response in I populations was restricted to the Bt strain used for selection: priming with heat-killed Bt1 (strain MTCC 6905) did not confer a survival advantage after a subsequent infection with live Bt1. Previous work with flour beetles (Roth *et al.* 2009; Khan *et al.* 2016) has revealed a similar degree of specificity of immune priming, allowing differentiation between strains of the same pathogen. Our results also show that this specificity extends to the pathogen strain used for priming, such that beetles primed with Bt1 do not gain a survival benefit against Bt. What molecular mechanisms underlie the evolved, strain-specific immune priming? Previous work suggests a role for phagocytosis (Pham *et al.* 2007; Weavers *et al.* 2016) or blood cell differentiation (Rodrigues *et al.* 2012) in mediating pathogen-specific immune priming in fruit flies and mosquitoes. Insects can also produce receptor diversity via alternative splicing of *Down syndrome cell adhesion molecule* to discriminate between different pathogens (Watson *et al.* 2005; Kurtz & Armitage 2006; Armitage *et al.* 2015). However, at the moment, the mechanism underlying evolved pathogen strain-specific immune priming in our experimental populations remains unclear. Recent work in vertebrate immunity suggests that cytolytic innate immune effectors such as natural killer cells also possess attributes of adaptive immune responses (Vivier *et al.* 2011) such that they can produce long-lasting, antigen-specific immune memory independent of B cells and T cells (O’Leary *et al.* 2006; Vivier *et al.* 2011). Hence, an additional possibility is the involvement of insect equivalents of natural killer cells in mediating strain-specific priming in our evolved lines. We also note that ancestral beetles already showed priming against lower doses of Bt infection (Khan *et al.* 2016), but not against the high dose of Bt that we used for selection. Since the cellular and molecular machinery for mounting a priming response was already present in the beetles, it was perhaps simply modified during experimental evolution to efficiently counter a higher pathogen dose. We thus speculate that the evolved priming response involved a quantitative (rather than qualitative) change –e.g. genes that were up-regulated in the ancestral population after a relatively low dose of Bt infection may be up-regulated further or faster in the evolved lines. In contrast to the high specificity of immune priming in I populations, we found that three of four PI populations showed generalized resistance against multiple strains of *B. thuringiensis*, suggesting that the divergent immune responses are driven by different mechanisms. Clearly, more work is needed to understand how specific immune memory and general resistance are achieved in insect immunity despite the lack of somatic recombination required for the development of adaptive immunity in vertebrates. We speculate that while vertebrate immune memory is mechanistically distinct, functionally it may not be as unique as traditionally believed.

One of the most striking outcomes of our experiment is the highly parallel yet mutually exclusive evolution of priming in I populations and increased resistance in PI populations. Priming evolved in only one PI population, only to be rapidly replaced by increased basal resistance. What prevented the evolution and maintenance of priming in PI populations? Conversely, why didn’t a resistance allele(s) sweep through I populations, despite the large potential selective advantage? One way to approach these questions is to determine the relative costs and benefits of hypothetical “priming” vs. “resistance” alleles, and ask whether their net benefit may vary as a function of I and PI treatments. It is clear that the survival benefit of evolved resistance was greater than the benefit of evolved priming, suggesting that a resistance allele should always outcompete a priming allele. Furthermore, the net benefit of evolved priming in I lines during experimental evolution was lower than that of evolved resistance in PI lines – after 11 generations, survival had increased marginally from 40% to 50% in I lines, but was as high as 80% in PI lines. Therefore, weakened selection for a resistance allele in I lines cannot explain the observed lack of resistance. What about the relative physiological costs of priming and resistance alleles? Generalized resistance in insects is associated with overexpression of fast acting non-specific immune effectors such as phenoloxidase and the production of reactive oxygen species (see Binggeli *et al.* 2014). Such immune effectors may facilitate rapid clearance of pathogens, but can also impose large physiological costs via damage to vital host organs such as Malpighian tubules (Sadd & Siva-Jothy 2006; Khan *et al.* 2017). In contrast, specific priming may require blood cell differentiation or Toll pathway activation (Pham *et al.* 2007; Rodrigues *et al.* 2012), which is potentially less toxic to host tissues (Hoffmann *et al.* 1996; Moret 2003; Rowley & Powell 2007). Thus, generalized resistance may be physiologically more costly than a specific priming response. McDade and colleagues have proposed a similar hypothesis for human immune function, drawing upon data from vertebrates to suggest that the cost of specific adaptive immune responses is lower than that of non-specific innate immune responses (McDade *et al.* 2016). Could a greater cost of basal resistance explain the pattern of evolution that we observe? A general mathematical model to explore the emergence of various forms of immune defense offers some clues (Mayer *et al.* 2016). Assuming that more effective defense incurs a greater maintenance cost, this model predicts that under frequent pathogen attack, a population’s growth rate is maximized by constitutively expressed resistance via innate immune responses. On the other hand, at a lower pathogen frequency, lower maintenance costs of inducible adaptive immunity make it more favorable. Our I and PI selection regimes broadly resemble these conditions of low vs. high frequency of infection: I populations received a single infection each generation, whereas PI populations were exposed to Bt antigens twice (though only one of them had live Bt). Thus, a large maintenance cost of generalized resistance combined with low frequency of infection in I populations may have prevented the evolution of resistance and favored the evolution of priming ability. These hypotheses need to be explicitly tested by identifying the immune effectors underlying specific vs. non-specific immunity, and evaluating their relative costs and benefits.

Finally, we note that post-infection survival may increase either via improved ability to kill pathogens or increased tolerance, where beetles do not kill pathogens more efficiently but reduce the cost of infection, immune response, or both (Schneider & Ayres 2008; Ayres & Schneider 2012). Since we estimated priming response and resistance using post-infection survival, it remains to be tested whether the broadly termed “resistance” involved direct pathogen clearance or improved tolerance. We were also unable to estimate the role of trans-generational immune priming, where parental exposure to pathogen can prime the immune state of their offspring (Zanchi *et al.* 2012; Khan *et al.* 2016). Hence, one possibility is that priming did occur in PI populations, but it was trans-generational priming (i.e. the survival benefit was transferred to offspring) rather than priming within the same generation. Thus, the stark increase in survival of PI beetles within the first generation of selection (from 40% to 68%) may have arisen from trans-generational immune priming, which would not be detected in our assays designed for within-generation priming. However, given that all experimental lines were standardized under relaxed selection for two generations, it is unlikely that trans-generational mechanisms could explain the observed difference in survival between PI and I populations.

In summary, we have documented the first experimental demonstration of rapid evolution of insect immune priming. Our work also represents a rare example where selection imposed by the same pathogen leads to divergent outcomes – either priming ability or basal resistance, but not both. We hope that our results will motivate further experiments to understand the dynamics and mechanistic basis of evolved priming vs. resistance. It is likely that an alternative vertebrate-like immune memory can evolve in invertebrates, but with different underlying molecules (Kurtz & Armitage 2006; Agaisse 2007). We also note that the general view of adaptive immunity as an exclusive mediator of immune memory in vertebrates has been recently challenged by the observation that resistance to re-infection can be achieved even without a functional adaptive immune system (Sun *et al.* 2009; Netea *et al.* 2016). Several studies are exploring the potential of the innate immune memory to aid novel therapeutic strategies for immunodeficiency and autoimmune disorders in vertebrates, including humans (reviewed in Netea *et al.* 2016). We suggest that insects are also a useful model system where both evolution and mechanistic basis of innate immune memory can be jointly studied.

## METHODS

### Beetle stocks

We generated an outbred *T. castaneum* stock population (ancestral, DA-IK) using adults from 10 wild-caught lines collected from different locations across India (Khan *et al.* 2016). We maintained this line as a large population (>5000 adults) on whole-wheat flour on a 45-day discrete generation cycle at 34 °C for 2 years before starting our experiments.

### Immune priming and challenge

We used the beetle pathogen *Bacillus thuringiensis* (DSM 2046) (Bt) isolated from a Mediterranean flour moth (Roth *et al.* 2009) to impose pathogen selection. For priming, we pricked adults between the head and thorax, using a 0.1mm insect pin (Fine Science Tools, CA) dipped in heat-killed bacterial slurry prepared from 10 ml freshly grown overnight culture of *B. thuringiensis* at 30°C (optical density of 0.95; adjusted to ~10^10^ cells in 100 μl insect Ringer solution) as described in Khan *et al.* (2016). Under natural conditions, individuals are more likely to experience live infection rather than killed pathogens. However, using live bacterial cells for priming would incur costs of infection and confound the effect of priming of the immune system by imposing mortality and selecting for priming as well as variants with greater basal resistance. To minimize the cost of and selection imposed by live infection, we used heat-killed bacteria that would prime the beetle immune system by eliciting an immune response without any cost of infection. We used insect Ringer (7.5 g NaCl, 0.35 g KCl, 0.21 g CaCl2 per liter) to perform mock priming. After 6 days, we challenged individuals with live bacterial culture (~10^10^ cells in 50 μl insect Ringer solution). This procedure delivers approximately 12000 live cells per beetle. Our previous work showed that basal level priming ability against a low dose of Bt (~8 × 10^3^ cells per beetle) already exists in this population, with primed beetles living significantly longer than unprimed controls after infection (Khan *et al.* 2016). However, when we tested beetles against a higher infection dose (~12 × 10^3^ cells per beetle), we did not find a significant difference in post-infection survival between primed and unprimed beetles (see Figure S1). Note that in both experiments we used the same dose of heat-killed Bt to prime beetles, and only the infection dose varied. We thus expected that the strength of priming would have to increase in order to counter the higher pathogen load used in the present study.

### Experimental evolution

For artificial selection, we used 4 selection regimes — control (C), primed only (P), infected only (I) and primed and infected (PI) (Figure 1A). We set up four replicate populations for each selection regime (C1 to C4, P1 to P4, I1 to I4, and PI1 to PI4). To initiate populations, we allowed ~1000 adults to oviposit in 500g of wheat flour. After 48h, we removed the adults and allowed offspring to develop for 3 weeks. We sexed offspring at the pupal stage and isolated males and females as virgins in wells of 96-well micro-plates (with ~25 mg wheat flour/well) for 2 weeks to allow adult emergence and sexual maturation. The pupal stage typically lasts for 3-4 days; hence we obtained 10-day-old adults for all experiments. During experimental evolution, we similarly collected pupae from each population to initiate the next generation. We primed 10-day-old adults from each population, either with heat-killed bacterial slurry (priming — P and PI regimes) or with sterile insect Ringer solution (mock priming — C and I regimes) as described above. For C and P regimes, we primed 100 males and 100 females per population; for I and PI regimes, we primed 200 males and 200 females per population since we expected higher mortality after subsequent infection with live bacteria.

After priming (or mock priming), we redistributed experimental beetles in wells of fresh 96-well micro-plates (Corning/Genetix) containing wheat flour. In all regimes, we found negligible mortality (<1%) after priming. After 6 days, we challenged individuals from I and PI regimes with live pathogen, whereas beetles from C and P regime were pricked with sterile insect ringer solution (mock challenge). Subsequently, we returned each individual to wells of fresh 96-well micro plates with flour. After two days, we counted the number of survivors from each population. For logistical reasons, we could not record survival for populations C4, P4, PI4 and I4 from generations 1-4 and for all replicate populations at generation 4. Next, we randomly selected 60 live individuals of each sex from each population and combined them in 500 g of wheat to mate and oviposit for 5 days. We then removed adults to allow offspring to develop. Note that although offspring number could vary across populations, we used excess flour (4 g/adult; experiments typically use less than 1g/adult; (Sokoloff 1977) to minimize resource competition among juveniles. The four replicates from each selection regime were handled on different days and maintained under continuous divergent pathogen selection for 11 generations.

At generations 8 and 11, we collected an additional set of pupae from each replicate population to generate standardized populations. These populations were maintained under relaxed selection (i.e. no mock injection, priming or pathogen infection) for two generations to minimize non-genetic parental effects (Faria *et al.* 2015; Gupta *et al.* 2016). For subsequent assays of evolved priming response and basal resistance (described below), we used individuals from these standardized populations.

### Quantifying evolved priming response and basal resistance

To measure evolved priming and resistance after 8 generations of selection, we randomly assigned 10-day-old virgin females from each standardized population to one of three treatments: (a) primed with heat killed cells of Bt (b) unprimed: pricked with insect Ringer and (c) uninfected control: pricked with insect Ringer (*n* = 16-24 females/treatment/replicate population/selection regime; Figure 1B). We then isolated them in the wells of 96-well micro-plates with access to flour. After 6 days, we challenged females from primed and unprimed treatments with live Bt1. Uninfected control females were pricked with insect Ringer. We noted survival of these standardized females derived after 8 generations of pathogen selection every 6 hours for 2 days and then every 24 hours for the following 50 days. We also noted survival of standardized males and females derived after 11 generations of pathogen selection every 6 hours for 2 days and then every 24 hours for the following 7 days (*n* = 16-26 sex/treatment/replicate population/selection regime). Using this protocol, we also re-evaluated the impact of bacterial infection and priming on females from the unhandled ancestral beetle population.

For each standardized population, we used Cox Proportional Hazard survival analysis to test beetle survival as a function of bacterial infection and priming. During experimental evolution, only eggs laid until a week after the immune challenge contributed to the next generation; hence, we expect selection to act strongly only during this period (selection window). We first analyzed beetle survival data until a week after immune challenge and considered individuals that were still alive at the end of the 7^th^ day as censored values. For standardized populations derived at generation 8, we also considered beetles that were still alive at the end of the 50^th^ day (full dataset) as censored values to estimate the delayed impact of pathogen selection. For each population, we calculated resistance to infection as the estimated hazard ratio of unprimed infected vs. uninfected control groups (Rate of death in unprimed infected group / Rate of death in the uninfected control group). A hazard ratio significantly greater than one indicates an enhanced risk of mortality in the infected group (i.e. lower resistance). In most populations, uninfected control beetles showed no mortality after 7 days (within the selection window) and the hazard ratio would be undefined. Hence, we introduced a single “death” in each of these uninfected control groups on the 7^th^ day, resulting in a more conservative but finite estimate of the hazard ratio. Finally, we calculated the survival benefit of priming as the hazard ratio of unprimed vs. primed groups. A hazard ratio significantly greater than one indicates increased risk of mortality in the unprimed group compared to primed individuals (i.e. a significant survival benefit of priming).

### Quantifying specificity of evolved immune priming and resistance

We used standardized females derived at generation 8 to test whether the evolved priming response and resistance was specific to the pathogen used for the selection experiment (i.e. Bt). To this end, we used a different pathogenic strain of *B. thuringiensis* (MTCC 6905) (Bt1) isolated from silkworm, to prime and challenge 10-day-old standardized virgin females as described in experiment 1. We monitored beetle survival every 6 hours for 2 days and daily around 10 pm for the following 7 days (*n* = 16-24 females/treatment/replicate population/selection regime). We analyzed survival data as described above. We were unable to estimate Bt1 priming in I4 lines for logistical reasons. In a separate experiment, we also tested whether the evolved priming ability (in I1, I2 and I4 lines) requires immune activation via priming with the same pathogen strain (homologous combination: primed and infected with Bt); or whether priming with a different strain could also induce the protective response (heterologous combination: primed with Bt1, infected with Bt). To do this, we primed 10-day-old standardized virgin females from C and I regime with heat-killed Bt and Bt1 as described above. After 6 days, we challenged each beetle with live Bt and noted its survival for 7 days (*n* = 16-24 females/treatment/replicate population/selection regime). We analyzed survival data using Cox proportional hazard analysis to compare beetle survival as a function of homologous priming (Bt-Bt) vs. heterologous priming (Bt1-Bt). We did not test population I3 because it did not show a priming response.

## ACKNOWLEDGEMENTS

We thank Aparna Agarwal, Rittik Deb and Saurabh Mahajan for critical comments on the manuscript, and NG Prasad for the *B. thuringiensis* strains. We acknowledge funding and support from a SERB-DST Young Investigator Grant to IK, a DST INSPIRE Faculty award to DA, and the National Centre for Biological Sciences (NCBS), India.

## AUTHOR CONTRIBUTIONS

IK conceived of experiments; IK and DA designed experiments; AP and IK carried out experiments; AP, IK and DA analyzed data; IK and DA wrote the manuscript with input from AP. All authors gave final approval for publication.

